# DNA Origami Presenting the Receptor Binding Domain of SARS-CoV-2 Elicit Robust Protective Immune Response

**DOI:** 10.1101/2022.08.02.502186

**Authors:** Esra Oktay, Farhang Alem, Keziah Hernandez, Aarthi Narayanan, Remi Veneziano

**Affiliations:** Department of Bioengineering, George Mason University, Fairfax, VA 22030, USA; National Center for Biodefense and Infectious Diseases, George Mason University, Manassas, VA 20110, USA; Institute for Advanced Biomedical Research, George Mason University, Manassas, VA 20110, USA

**Keywords:** SARS-CoV-2, Vaccine, DNA Origami, Receptor Binding Domain, Nanoparticle Vaccine, CpG Oligodeoxynucleotide, COVID-19

## Abstract

Effective and safe vaccines are invaluable tools in the arsenal to fight infectious diseases. The rapid spreading of severe acute respiratory syndrome coronavirus 2 (SARS-CoV-2) responsible of the coronavirus disease 2019 pandemic has highlighted the need to develop methods for rapid and efficient vaccine development. DNA origami nanoparticles (DNA-NPs) presenting multiple antigens in prescribed nanoscale patterns have recently emerged as a safe, efficient, and easily scalable alternative for rational design of vaccines. Here, we are leveraging the unique properties of these DNA-NPs and demonstrate that precisely patterning ten copies of a reconstituted trimer of the receptor binding domain (RBD) of SARS-CoV-2 along with CpG adjuvants on the DNA-NPs is able to elicit a robust protective immunity against SARS-CoV-2 in a mouse model. Our results demonstrate the potential of our DNA-NP-based approach for developing safe and effective nanovaccines against infectious diseases.

## Introduction

With the emergence of severe acute respiratory syndrome coronavirus-2 (SARS-CoV-2) and thus coronavirus disease 2019 (COVID-19), our healthcare systems have been facing unprecedented challenges.^1^ The COVID-19 pandemic has already resulted in the death of millions around the world and forced authorities to impose mandatory quarantines to limit viral spreading.^2^ SARS-CoV-2 is an emerging and highly infectious RNA virus that belongs to the β-coronavirus genus, which contains other viruses that are responsible for previous major outbreaks.^3^ One key strategy to limit the spreading of SARS-CoV-2 is the development of safe and effective vaccines that can provide long-lasting immunity and protect against all variants.^4,5^ Current vaccines strategies primarily use all or part of the viral spike protein (S) as an immunogen that can be delivered in various forms (e.g., mRNA, DNA plasmid, and protein).^6^ Among all of the candidate vaccines that reached clinical trials, only three have been issued by FDA for Emergency Use Authorization. Two are mRNA-based vaccines [the mRNA-1273 from Moderna and the BNT162b2 from Pfizer/BioNTech (COMIRNATY®)] with a >90% efficacy reported and one is a viral-vector-based vaccine [Ad26.CoV2.S from Johnson & Johnson (Janssen)] with 66% efficacy.^7–9^ Numerous traditional vaccine strategies that rely on either killed-inactivated or live-attenuated viruses are currently under development, some of which are in Phase III of clinical trials, but strikingly none have yet received FDA approval.^10^

Other methods such as subunit vaccines are currently being developed, as they represent a safe way to deliver antigens. However, monovalent antigens are known to often trigger low immunogenicity that results in limited protection, which can be mitigated by displaying them in multivalent form on nanoparticle (NP) carriers.^11,12^ Because NP-based vaccines do not use viral genetic materials and are assembled with biocompatible materials,^13^ they also reduce safety concerns.^14^ Additionally, their chemical nature, surface compositions, and modifications can improve vaccine stability, thereby helping to prolong their shelf-life, extend their bioavailability, and facilitate their cellular uptake.^14,15^ To this day, more than 60 NP-based SARS-CoV-2 vaccines have been developed, showing the high interest for these strategies.^16^ Among these candidates, most of them deliver the receptor binding domain (RBD) of the spike S1 protein or of the full spike protein.^17,18^ The RBD appears to be a potent immunogen and one of the main targets of neutralizing antibodies found in vaccinated and infected people.^19,20^ This domain is also crucial during viral infection due to its role in interacting with host cells to promote viral entry. The RBD is recognized by the angiotensin converting enzyme 2 (ACE2) receptor and therefore plays a vital role in viral tropism and spreading to cells expressing ACE2 receptors.^21^ The structural feature of the RBD has been identified as having either closed or open states.^22^ In the closed conformation, three RBDs of the trimeric S protein can mask themselves to avoid recognition by the immune system. The S protein undergoes conformational changes and only one RBD reveals its binding motif to interact with ACE2 in open conformation, which is then followed by the sequential opening of the other two RBD motifs and thereby forms the trimeric S protein-ACE2 receptor complex.^22–24^ Many studies have shown that a multivalent display of RBDs induces a stronger immune response and a higher level of neutralizing antibody titer in comparison with monomeric RBD.^25–28^ However, the limited control offered by NPs does not allow for assessing the role of structural parameters related to antigen presentation (e.g., density, stoichiometry, nanoscale organization, and NP geometries) on cellular uptake and immunogenicity that could enable rational design of vaccines.^29,30^

Scaffolded DNA origami nanoparticles (DNA-NPs), on the other hand, provide an ideal biocompatible platform for assessing antigen presentation parameters.^31,32^ Indeed, DNA-NPs can be designed in any geometry and size^32,33^ while permitting the patterning of biomolecules with nanoscale precision, which allows for the modulation of antigen stoichiometry and facilitates multiplexing^34,35^. Recently, DNA-NPs have been used to present antigens in controlled stoichiometry and nanoscale organization to modulate the activation of immune cells. Specifically, Veneziano et al. showed that presenting HIV-glycoprotein (eOD-GT8) antigens on DNA-NPs in various nanoscale organization and stoichiometry led to the modulation of B-cell activation by inducing the clustering of B cell receptors *in vitro*.^34^ Thus, using DNA origami could inform the rational design of vaccines and represent a promising alternative nanocarrier for effectively and safely delivering viral antigens. In this study, we developed a DNA-NP vaccine platform (DNA-NP nanovaccine) that can accommodate multiple copies of a single immunogen and adjuvant simultaneously. We also assessed its immunogenicity and efficacy *in vivo*. Specifically, we built a DNA pentagonal bipyramid that presented the RBD antigen in a trimeric form, along with CpG ODN 1018 (cytosine-phosphorothioate-guanine oligodeoxynucleotides) adjuvant, to assess efficiency of the DNA-NP nanovaccine in mounting a strong and protective immune response (Fig. 1).

**Fig. 1.**
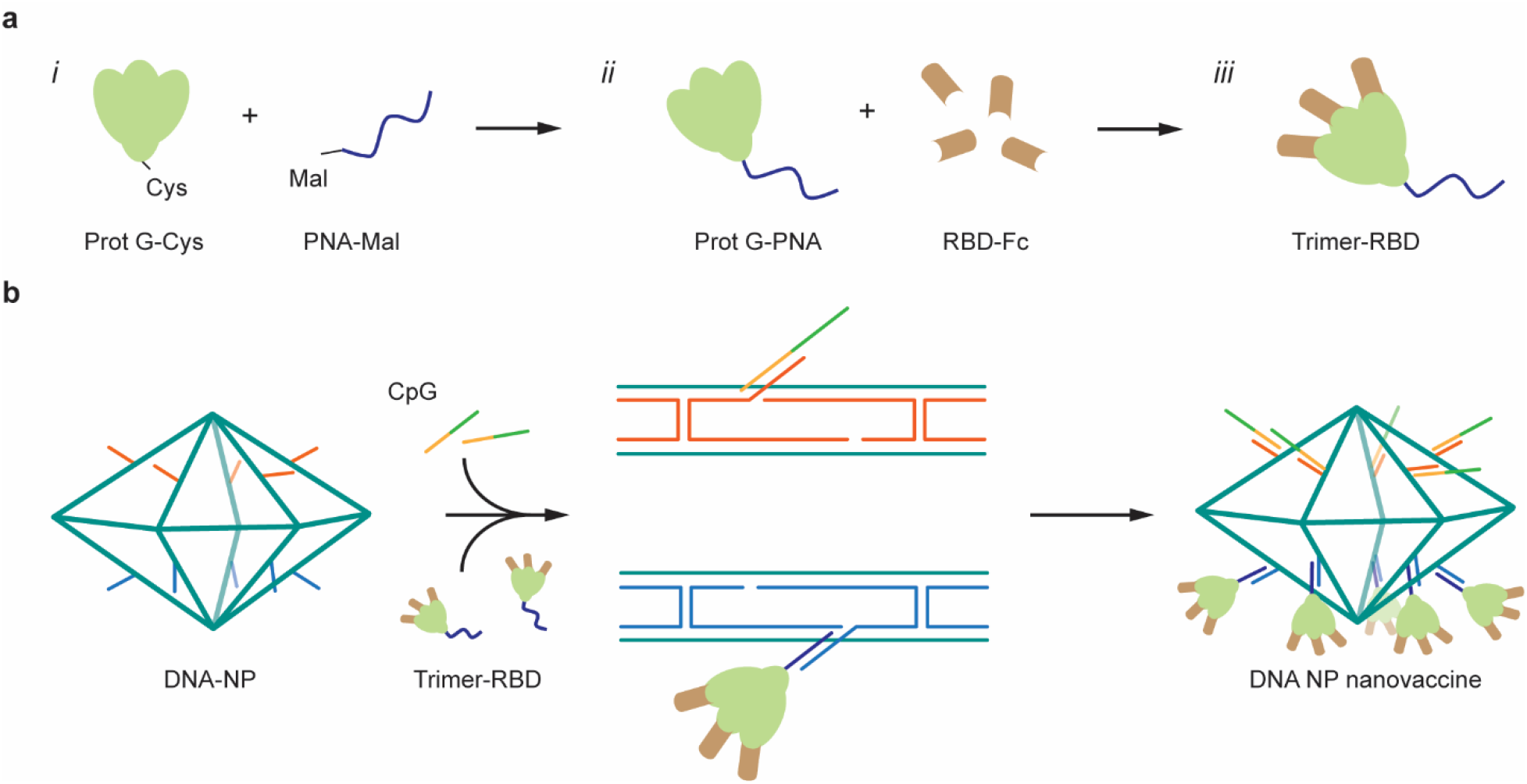
Assembly of the DNA-NP nanovaccine. **a** Formation of the peptide nucleic acid (PNA)-RBD trimer by coupling three RBD-Fc on the three Fc-binding domains of a protein G (PG): ***i***. A PNA strand is conjugated to a PG via maleimide chemistry. ***ii***. PG-PNA is used to couple three RBD-Fc. ***iii***. The trimer is purified from the free RBD-Fc. **b** Two separate faces of the PB are modified with ssDNA overhangs on defined locations to facilitate attachment of the PG-RBD complex and CpG adjuvants.

## Results

### Constructing 3D-wireframe DNA origami nanoparticles

Using the DAEDALUS (DNA Origami Sequence Design Algorithm for User-defined Structures) software^33^, we designed a DNA pentagonal bipyramid (PB) (Supplementary Fig. 1) with a diameter of 36.5 nm (52 base pairs [bps] edge length). The sub-100 nm size of our NP was chosen to facilitate lymph drainage and promote uptake by antigen presenting cells, as previously demonstrated in the literature regarding the optimal size for efficiently draining to lymph nodes and current nanoparticle-based approaches such as protein and lipid nanoparticles.^36–38^ The PB was assembled with a long single-stranded DNA (ssDNA) scaffold that was produced by asymmetric PCR^39,40^ (Supplementary Tables 1 and 2 and Supplementary Fig. 2) and folded with 44 staple strands (sequences of scaffold and staple strands are listed in Supplementary Table 1 and Supplementary Tables 3 to 5, respectively). The PB edges were designed to display up to 10 ssDNA overhangs per face (2 overhangs per edges) at the 3’-end of selected staple strands, in addition to 10 ssDNA overhangs on the side edges (Supplementary Fig. 3) to anchor antigens and CpG adjuvants modified with complementary overhangs via direct hybridization. These overhangs were designed to be orthogonal to each other in order to facilitate asymmetric modification of the NPs, thus enabling simultaneous presentation of the antigens on one face and delivery of the conjugated adjuvants on the other face. The correct folding of the PB constructs was validated with agarose gel electrophoresis and dynamic light scattering (DLS), which showed a well-folded monodisperse particle population with a folding yield estimated at 96% (Fig. 2a and Supplementary Fig. 4). The diameter of the PB was measured with DLS at 51.4 nm for a theoretical diameter of 46.4 nm diameter for particles including overhangs. This result is consistent with previous results reported with this type of NPs.^33,34^

**Fig. 2.**
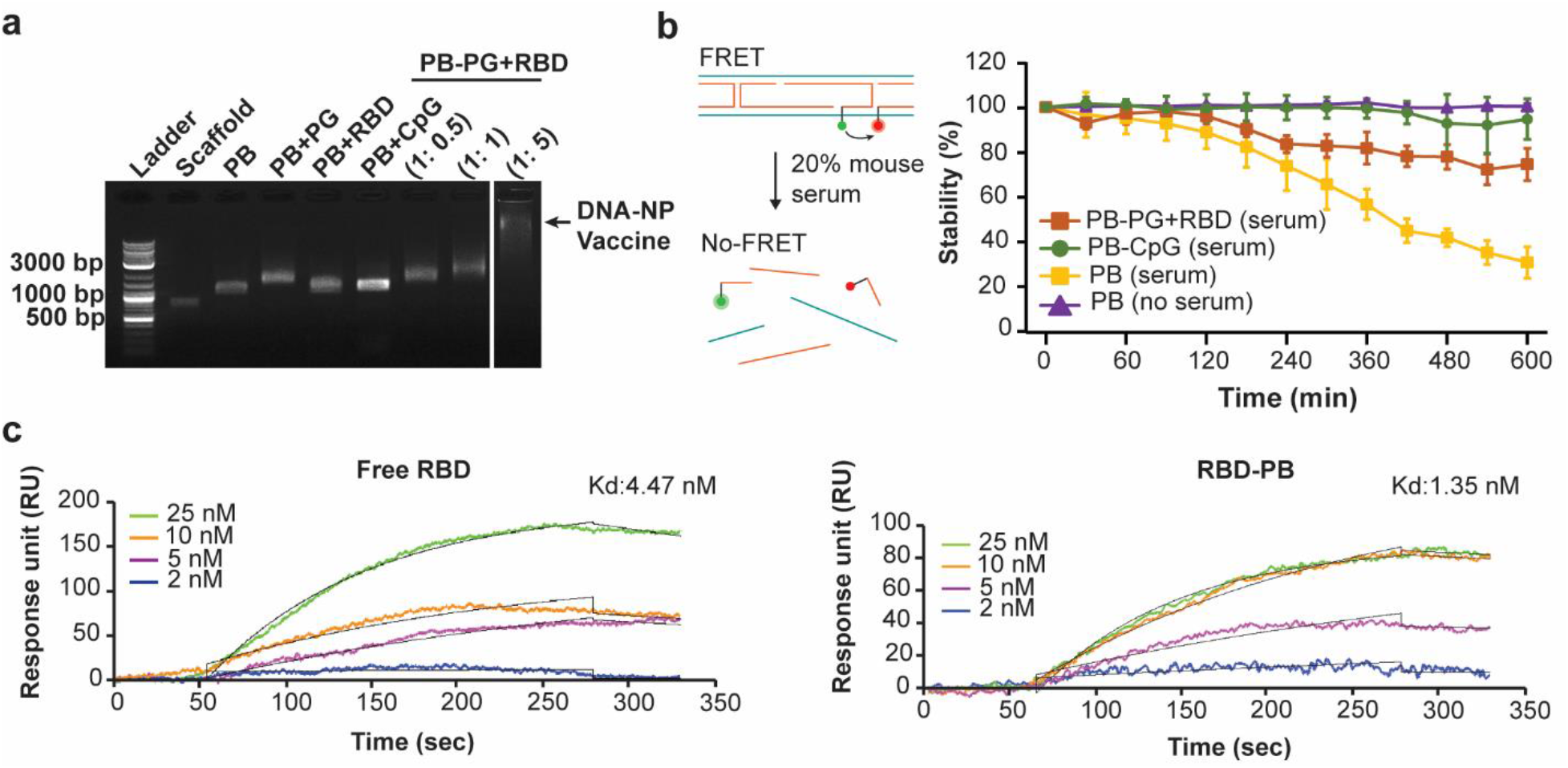
Characterization of PB DNA-NP nanovaccines. **a** CpG hybridization and RBD attachment to DNA origami nanoparticle. To evaluate binding of PG and RBD, PG-PNA and RBD-Fc were first mixed in different molar concentrations (1:0.5, 1:1, 1:5). Then, DNA-NP was added into the reaction solution (the molar ratio between DNA-NP and PG is 1:2). Conjugation of PG-RBD to DNA-NP was evaluated via 1% agarose gel electrophoresis. **b** Determination of nanovaccines stability via an *in vitro* FRET-based assay. Nanoconstructs were incubated for 10 hours in 20% mouse serum and the changes in the FRET efficiency were examined to determine stability. PB-CpG and RBD-PB-CpG showed good stability over the entire 10-hour period. The fluorescence intensity of bare PB decreased approximately 60% after 10 hours. (Yellow line: PB positive control, N=3 samples; violet line: PB negative control, N=3 samples; PB-CpG, N=4 samples; RBD-PB-CpG, N=2 samples). Data were shown as mean ± SD on graph. **c** Representative binding kinetics measurement via surface plasmon resonance. Four different concentrations (25 nM, 10 nM, 5 nM, and 2 nM) of free RBD monomer and trimeric RBD-PB were tested for their binding kinetics flowing over immobilized ACE2 receptor. Measured Kd between free RBD monomer and ACE2 was determined at 4.47 nM, and Kd between RBD-PB and ACE2 receptor was determined at 1.35 nM.

### Reconstitution of trimeric-RBD via Protein G-Fc conjugation strategy

Next, we prepared the reconstituted RBD-trimer that we used as an immunogen to display on the DNA-NPs. We used a commercially available modified version of the protein G (cys-PG) that contains a single cysteine residue on its N-terminal. A peptide nucleic acid strand (PNA) was conjugated to the cys-PG via a maleimide group (Fig. 1a). PNA is a nucleic acid analog composed of peptide bonds and nucleobases capable of hybridizing DNA via Watson Crick base-pairing with a higher affinity than DNA:DNA hybridization.^41^ The PG-PNA conjugation was validated with sodium dodecyl sulphate polyacrylamide gel electrophoresis (SDS-PAGE) (Supplementary Fig. 5) and then purified with centrifugal filtration. The concentration of PG was estimated based on the concentration of PNA. We then reacted various ratios of RBD-Fc and PG-PNA to ensure complete reconstitution of the RBD trimer and validated their formation with native and SDS-PAGE (Supplementary Fig. 6).

### DNA origami nanovaccine presenting CpGs and RBD-trimers

To control the organization and stoichiometry of the RBD-trimers on the surface of the DNA-NPs, we hybridized 10 copies of PG-PNA to specific overhangs displayed on the surface of the DNA-NPs (Fig. 1b). The RBD-trimer was then assembled on the antigen presenting face of the PB through PG-PNA, and 10 CpG strands were also hybridized on the opposite face of the PB via hybridization on a second set of overhangs (Fig. 1b). We used the CpG ODN 1018 (cytosine-phosphorothioate-guanine oligodeoxynucleotides), which contains a fully phophorothioated backbone and has been already proven to be efficient in an FDA-approved Hepatitis B vaccine.^42^ Specifically, CpG 1018 is known to stimulate Th1-biased CD4^+^ T cells characterized with pro-inflammatory cytokine IFN-γ secretion as a part of antiviral immunity.^43,44^ After validating attachment of PG-PNA to specific overhangs via PAGE (Supplementary Fig. 7), we confirmed the attachment of the protein complex (RBD-PG) to PB NP, along with CpG hybridization, via agarose gel electrophoresis (Fig. 2a).

### Characterization of the stability of DNA nanoparticles

Prior to assessing the efficacy of our DNA-NP nanovaccines *in vivo*, we evaluated their stability in simulated physiological conditions. We used a fluorescent resonance energy transfer (FRET)-based assay with two FRET reporter pairs (Supplementary Table 6) located on two different edges of the PB (Fig. 2b). The sites selected do not interfere with the antigen or the CpG binding sites. We used fluorescein (FAM) and tetramethylrhodamine (TAMRA) as donor and acceptor dyes, respectively. The FRET pair were designed with a distance of 8 bases (∼3 nm) between dyes to maximize the FRET efficiency. The stability of NPs was evaluated throughout a 10-hour incubation in PBS containing 20% mouse serum, and the changes in fluorescence intensity of the acceptor dye was used to monitor the degradation rate (Fig. 2b). The stability of the NPs was measured from the FRET efficiency as calculated in Wei et al. (2013).^45^ Our FRET results showed that 40% of the Bare PB was still intact after 10 hours. Interestingly, PB modified with CpG strands, as well as with a combination of RBD, showed significantly increased stability with more than 70% of particles still intact after 10 hours. These results demonstrate that coating DNA-NPs with protein and modified nucleic acids can participate in increasing the stability against nucleases without the need for complex chemical modifications as seen in extant literature.^46^

### Comparing binding affinity of free RBD monomers and trimeric RBD on DNA NP

Next, we used surface plasmon resonance (SPR) to validate the accessibility of the antigens on the surface of the NPs by evaluating the binding kinetics of our DNA-NP nanovaccines presenting RBD-trimer in comparison with the free RBD. We modified the SPR gold sensor surfaces with the soluble ACE2 receptor. Immobilization of ACE2 was consistent across all the experiments performed with an average of 1540.3 ± 292.2 (mean ± SD) RU (Supplementary Fig. 8). Soluble RBD antigens at various concentration (2 nM, 5 nM, 10 nM, and 25 nM) were injected over the ACE2 receptor and the equilibrium dissociation constant (Kd) between ACE2 and the free RBD was calculated at 4.47 nM (Fig. 2c). We used the DNA-NP nanovaccines presenting the RBD-trimer at different concentrations (2 nM, 5 nM, 10 nM, and 25 nM equivalent RBD concentration) and determined a Kd of 1.35 nM (Fig. 2c). In each SPR experiment, a single- cycle kinetic measurement was performed, without regeneration of the surface, and with a long dissociation time before each injection. (Supplementary Fig. 9) Table 1 summarizes the values (mean ± SD) associated to the binding association/dissociation rate constants (kon and koff, respectively) and the Kd calculated from the SPR curves. The Kd values determined in our studies are in the same range as previously published studies (Supplementary Table 7). ^47–49^

**Table 1.**
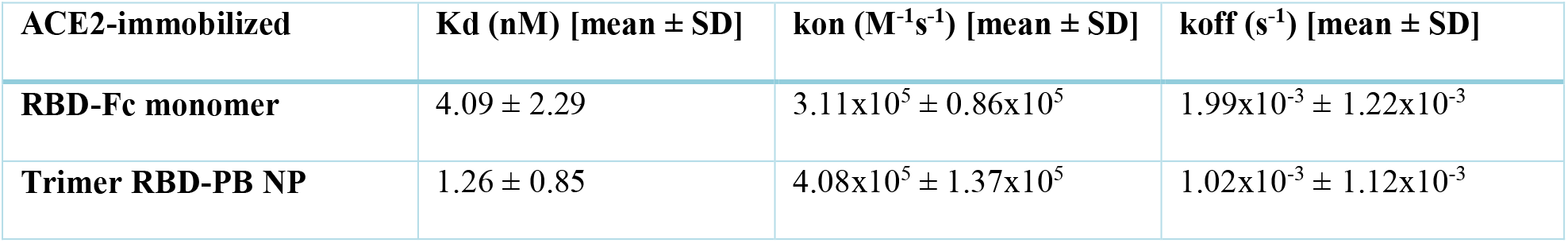
Binding kinetic measurements of soluble RBD and RBD-PB NP on SPR sensor chips grafted with the ACE2 receptor.

### Immunizing BALB/c mice with DNA origami nanovaccines elicits high antibody response

After validating the availability of the RBD domain on the DNA-NPs, we prepared four different constructs to perform immunization assays with a BALB-C mouse model. Bare PB samples were prepared as a negative control along with RBD-PB and RBD-PB-CpG as candidate vaccines. Recent studies, including ongoing clinical trials with RBD, have used doses of RBD ranging from 1 to 90 μg.^50–53^ We assessed two different RBD quantities (1 μg and 5 μg) presented on our PB. The quantity of RBD delivered was increased by injecting a higher concentration of NPs per dose. Two doses of the constructs were injected intramuscularly (IM) to the mice with a three-week interval (Fig. 3a). Six weeks after the first injection, mice were euthanized, and the serum was collected to perform further tests.

**Fig. 3.**
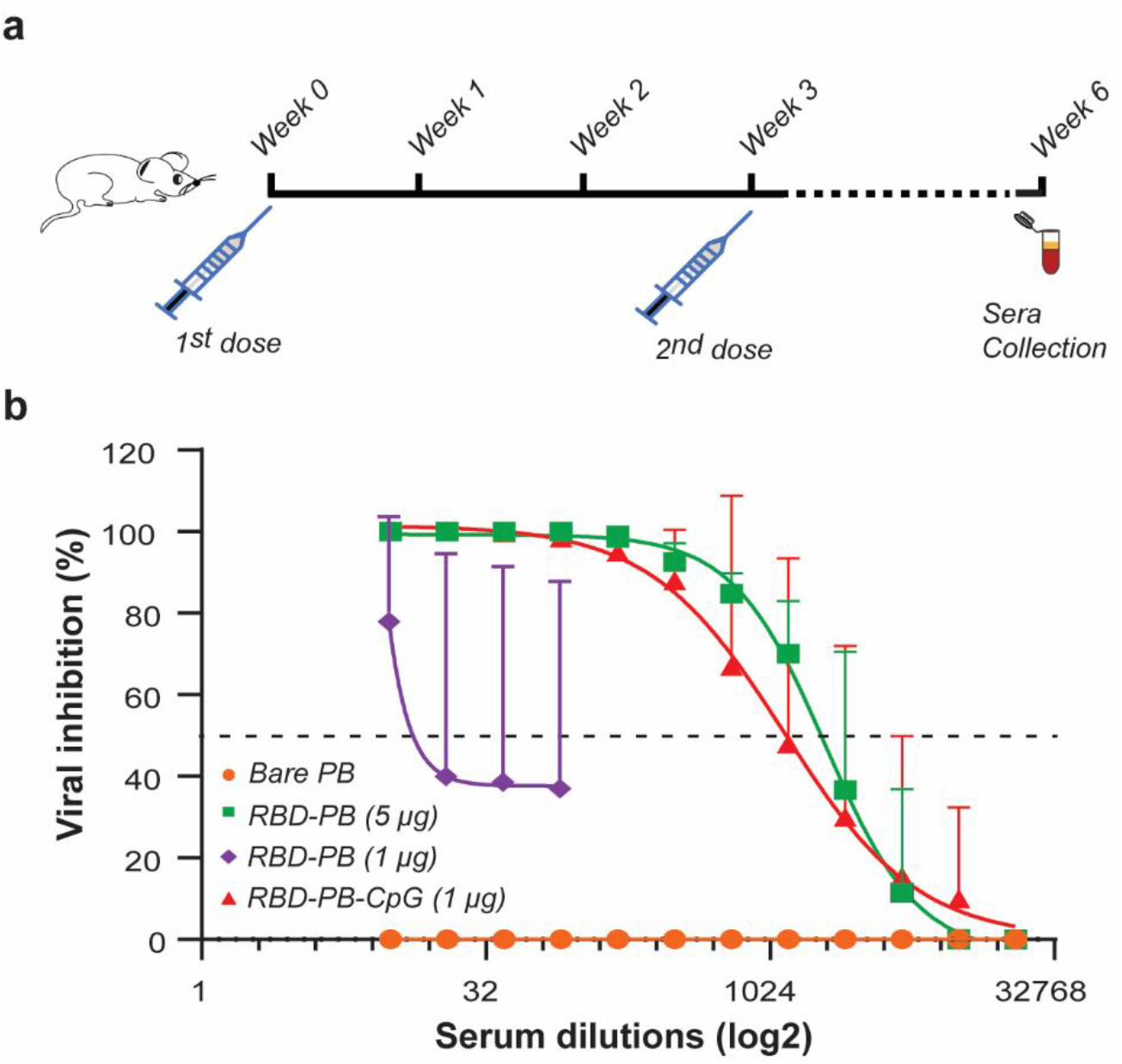
Immunization studies performed with four different PB DNA-NP nanovaccine constructs. **a** Two doses of PB DNA-NP nanovaccines (IM injection; 50 μl each) were administered with a three-week interval. Sera were collected at the end of the 6^th^ week to assess antibody titers via PRNT and end-point dilution assays. **b** Among four different animals (N=5 per study group), significant viral inhibition was observed in mice immunized with RBD-PB (1 μg), RBD-PB (5 μg), and RBD-PB-CpG (1 μg). Data were shown as mean ± SD on graph. Two-way ANOVA was performed to evaluate the statistical significance of data (p<0.001).

The presence and neutralizing efficacy of antibodies in immunized serum samples obtained from vaccinated animals were determined using a plaque reduction neutralization assay (PRNT) carried out by conventional methods.^54,55^ Our PRNT results demonstrate that PB with a 5-μg dose of RBD, 1-μg dose of RBD plus CpG ODN 1018 were all highly effective in eliciting a strong neutralizing antibody response. The data showed strong inhibition of viral infectivity from the vero monolayer at >1:1024 dilution, whereas PB alone had no effect on viral inhibition and PB with 1 μg RBD had limited efficacy (Fig. 3b).

### DNA origami nanovaccines protect against aerosol challenge with live SARS-CoV-2

The initial assessment of immunogenicity also suggested that no adverse events were observed in the vaccinated animals under the dosage regimen employed. As a next step, we evaluated if the immunization regimen could confer protection in the face of a lethal challenge by SARS-CoV-2 in the K18-ACE2 mouse model (transgenic mice expressing the ACE2 receptor) ^56–58^. We injected five of them with five different DNA-NP nanovaccine constructs, namely Bare PB, RBD-PB (1 and 5 μg), RBD-PB-CpG (1 μg), and PB-CpG. After injecting two doses of the PB vaccine constructs with a three-week interval, the mice were exposed to the live SARS-CoV-2 (Isolate Italy, INMI1)^59^ via intranasal route at a dose of 5×10^4^ plaque-forming unit (pfu) per mouse (Fig. 4a). The survival rate of each animal was evaluated for 14 days, along with daily monitoring of their weight loss (Fig. 4b and 4c). At Day 7 after viral exposure, mice immunized with Bare PB, RBD-PB (1 μg), and PB-CpG showed a nearly 30% weight loss, whereas mice immunized with RBD-PB (5 μg) and RBD-PB-CpG (1 μg) did not show weight loss. At Day 8, the groups injected with Bare PB, RBD-PB (1 μg), and PB-CpG had a drop in the survival rate with 60% mortality for PB-CpG and 80% mortality for Bare PB and RBD-PB (1 μg). At Day 14, animals vaccinated with 5 μg dose of RBD-PB showed only 40% mortality. At Day 14, only two mice injected with PB and PB-CpG survived. Interestingly, the PB DNA-NP nanovaccine construct prepared with CpG plus 1 μg dose of RBD showed no mortality and also no body weight loss over the 14-day period, thus confirming the protection provided by our nanovaccine construct. Recent studies have shown that a CpG 1018 adjuvant is effective in the induction of neutralizing antibodies and Th1-biased cell responses against SARS-CoV-2, which seems to be confirmed in our study.^43,44,60^ According to our results, administering PB with CpG and 1 μg of RBD together enhanced immunity against the virus. Furthermore, using CpG clearly improved immunization and allowed the use of lower antigen quantities to trigger a specific and strong immune response.

**Fig. 4.**
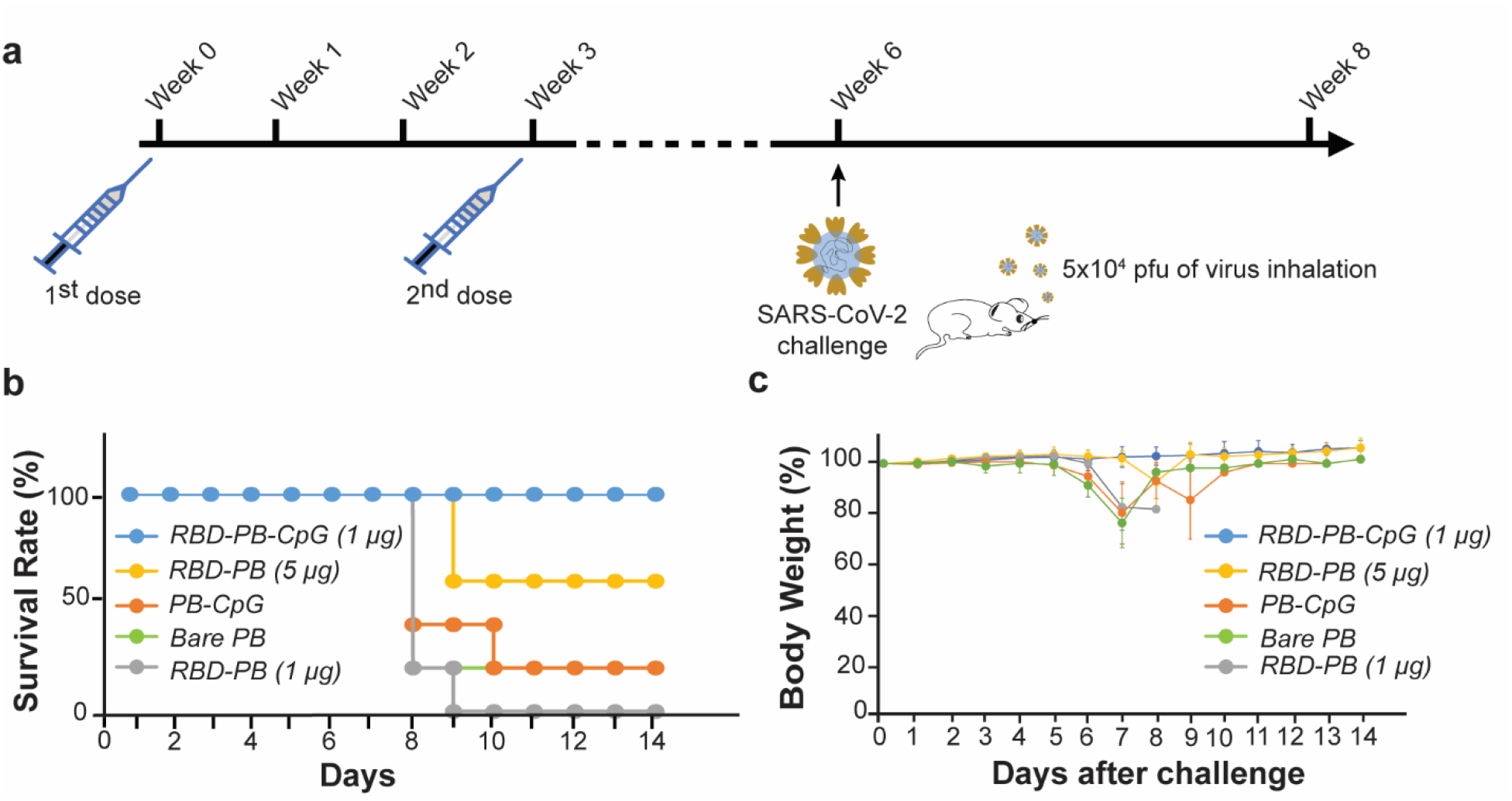
Challenge studies with SARS-CoV-2 virus. **a** Animals (n=5 per study group) immunized with two doses of PB DNA-NP nanovaccine constructs were subjected to challenge with SARS-CoV-2 via intranasal route of infection at Week 6. **b** Body weight changes (mean ±SD) and **c** survival rate were monitored for 14 days after intranasal viral challenge. After 14 days, only one animal survived among mice vaccinated with bare PB and one animal vaccinated with PB-CpG. All mice vaccinated with RBD-PB (1 μg) succumbed at Day 8. Three mice survived among the animals vaccinated with RBD-PB (5 μg). There was no weight loss and death observed among mice vaccinated with (1 μg) RBD-PB-CpG. **c** Weight loss of post-challenged mice were compared with one-way ANOVA, Tukey, and Kruskal-Wallis post-hoc test (p<0.001).

## Conclusion

In this study, we designed a 3D DNA PB and displayed 10 copies of a SARS-CoV-2 RBD trimer along with 10 CpG adjuvants on the two opposite faces of the NP. The viral challenge experiments showed that even small antigen doses delivered by the DNA-NPs are sufficient to provide protection against the virus when administered along with adjuvants. These results clearly demonstrate the potential of this strategy to develop a vaccine and highlight the importance of testing the spatial organization and stoichiometry of antigens to maximize the cellular response. These DNA-NPs could be used to present multiple types of immunogens simultaneously to develop a ‘broad spectrum’ vaccine targeting multiple viral strains.

## Methods

### Reagents, consummables and kits

The oligonucleotides used as primers in the asymmetric polymerase chain reaction (aPCR) and as staple strands for the folding of nanoparticles, along with the CpG ODN 1018 adjuvant were purchased from Integrated DNA Technologies (IDT). Fluorescently labelled oligonucleotides used in FRET assay, FAM (donor) and TAMRA (acceptor), were also procured by IDT. The deoxynucleotide triphosphates (DNTPs) mix (cat. no: N0447L), the Quick Load Purple 1 kb plus DNA ladder (cat. no: N0550S), and the M13mp18 circular single stranded DNA template (cat. no: N04040S) was obtained from New England Biolabs (NEB). The AccuStart Taq DNA polymerase HiFi enzyme was obtained from Quanta Biosciences. Low melt agarose (cat no: 89133-104) was provided by IBI scientific. The Zymoclean Gel DNA recovery kit was provided by Zymo Research (cat. no: D4008). The 10 kDA MWCO (cat. no: UFC5010) and 100 kDA MWCO (cat no: UFC5100) AmiconUltra 0.5 centrifugal filters, the 11-mercaptoundecanoic acid (CAS no: 71310-21-9) and L-Tryptophan (CAS no: 73-22-3) were obtained from Sigma Aldrich. The RBD, Fc (cat. no: SPD-C5255) and the biotinylated human ACE2/ACEH, His, Avitag (cat. no: AC2-H82E6) were provided by ACRO Biosystems. The cys-protein G was purchased from Prospec (cat. no: pro-1238). The PNA-Maleimide was procured by PNA Bio. The Gold sensor chips for SPR experiments were obtained from Nicoya Lifesciences (cat. no: SEN-AU-100-10). Greiner 384-well polystyrene flat bottom microplate was purchased from Cellvis (cat no: P384-1.5H-N). PageBlue Protein Staining Solution (cat. no: 24620), 1% crystal violet (cat. no: C581-25) and 20% ethanol solution (cat. no: BP2818-4), formaldehyde (cat. no: F79p-4) and 0.6% agarose (cat. no: 16500100) were purchased from Thermo Fischer Scientific. 1 mM sodium pyruvate was purchased from VWR (cat. no: 45000-710).

### Cell lines and animals

Vero cells were purchased from ATCC. BALB-C mice were purchased from Jackson Laboratories. K18-Ace2 (B6.Cg-Tg(K18-ACE2)2Prlmn/J) mice were purchased from Jackson Laboratory (Stock no: 034860).

### Preparation of the DNA origami nanoparticle nanovaccine

#### Pentagonal bipyramid (PB) scaffold production

The PB single stranded DNA (ssDNA) scaffold was synthesized via aPCR using the protocol described in Veneziano et. al. in 2018.^61^ Briefly, 50 μl reaction mix was prepared with 25 ng of M13mp18 template, the PB primer set with a 1:50 molar ratio (1 μM forward primer, 20 nM reverse primer), 1X HiFi buffer (provided by Quanta Biosciences) supplemented with 2 mM of Magnesium Sulfate (MgSO4), 0.2 mM of the dNTPs mix, and 1.25 U of AccuStart Taq DNA Polymerase HiFi. The aPCR cycle were performed in a Bio-Rad T100 Thermal Cycler as follow: activation at 94°C for 1 min followed by 30-40 cycles of 94°C for 20 s, 55°C for 30s, and the amplification step at 68°C for 2 min. Low melt agarose (1.2%) preloaded with Ethidium bromide were used to visualize and purify the ssDNA scaffolds. The ssDNA band were cut from the gel and purified using the Zymoclean Gel DNA recovery kit following the vendor instructions. The ssDNA scaffold was quantified using Nanodrop and the purity was assessed via gel electrophoresis.

#### Folding of PB nanoparticle (NP)

The ssDNA scaffold was folded into PB NP using a 1:10 molar ratio of scaffold *vs* staple strands. The folding reaction was performed via an overnight annealing in TAE-Mg^2+^ buffer (40 mM Tris, 20 mM acetic acid, 2 mM EDTA, and 12 mM MgCl_2_, pH 8.0) from 95°C to 4°C as previously described.^33^ PB folding was validated via agarose gel electrophoresis and the yield was quantified using Image J.

#### Conjugation of PNA strand to the Cys-Protein G (PG)-Cysteine

PG with N-terminal cysteine was reduced with 10-fold molar excess of tris(2-carboxyethyl)phosphine (TCEP) at room temperature for 15 min. After 15 min, PG was filtered via 10 kDa AmiconUltra-0.5 centrifugal filter to remove excess of TCEP and reacted overnight with 3-fold molar excess of PNA-Maleimide at 4°C. PG-PNA purification was achieved with 11 filtration steps using 10 kDa AmiconUltra-0.5 centrifugal filter to remove the unbound PNA. [The sequence of PNA with maleimide is: ‘SMCC-GGK-cagtccagt-K’, which is composed of amino acid residues (shown as uppercase) and bases (shown as lowercase)].

#### Reconstitution of the RBD trimer immunogen

Purified PG-PNA was mixed with 5-fold molar excess of RBD-Fc and incubated in 1X phosphate buffer saline (PBS) at 37°C for 1.5 hr and purified with AmiconUltra filter to remove the excess of monovalent RBD-Fc.

#### Functionalization of the PB nanoparticle with the RBD trimers and the CpG strands

Attachment of RBD-PG-PNA to PB NP was performed via hybridization to DNA overhangs displayed by staple strands on the DNA NP at 37°C for 1.5 hr. The molar ratio between PB NP and RBD-PG-PNA was 1:2, respectively. For conjugation with CpG, a 10-fold molar excess of CpG ODN 1018 was used and the reaction was done at 37°C for 1.5 hr. Reaction product was filtered from excess amount of CpG and PG-RBD protein complex using centrifugal filter. Purified RBD-PB-CpG was kept at 4°C before further use. (Sequence of CpG 1018 with linker complementary to specific overhangs on PB is the following: 5’-T*G*A*C*T*G*T*G*A*A*C*G*T*T*C*G*A*G*A*T*G*A<u>ACTTCATGGTCCTAACTT</u>-3’. (* indicate phosphorothioate linkage and the underlined sequence indicates the linker sequence for hybridization to the overhangs).

### Characterization of DNA origami nanoparticle nanovaccine

#### Fluorescence resonance energy transfer (FRET)-based stability assay

Folded PB (bare PB, PB-CpG, and RBD-PB-CpG) were modified with two FRET pairs (Donor: fluorescein [FAM] and acceptor: TAMRA). The FRET NPs (80 nM) were incubated in 20% (v/v) mouse serum to assess the relative stability ratio throughout a 10-hour period. The fluorescence measurements were performed in a Microplate reader (Tecan Safire2) with an excitation at a wavelength of 455 nm (20 nm bandwidth) and the emission spectra collected from 500 nm to 700 nm (20 nm bandwidth). The samples were loaded into Greiner 384-well polystyrene flat bottom black microplate. The following equations were used to calculate the FRET efficiency and determine the rate of degradation. We used the change in the fluorescence intensity of donor dye over time according to the method proposed by Wei et. al..^45^ According to this formula,

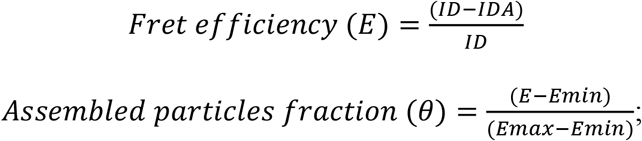

where ID represents the intensity of donor dye from only donor bearing PB NPs and IDA represents the intensity of donor dye from donor-acceptor bearing PB NPs.

#### Binding kinetics measurement of the RBD on ACE2 receptors via surface plasmon resonance

Binding association and dissociation kinetics between ACE2 and RBD-Fc or PB-RBD were measured with an Open Surface Plasmon Resonance (Open SPR) instrument provided by Nicoya Lifesciences. Gold sensor chips were first prepared by immersion in 11-mercaptoundecanoic acid and kept at room temperature for 48 hrs. Prior to the SPR measurements, the amino gold surface was biotinylated by incubation with 100 μl EDC/NHS mix from a stock of 200 mM EDC and 50 mM NHS for 3 min at room temperature, which was followed after thorough rinsing with water by an incubation with BSA-biotinylated (0.5 mg/ml) in 10 mM HEPES, 150 mM NaCl buffer at pH 7.2 for another 3 min at room temperature. The sensor chip was then loaded in the SPR device and a streptavidin solution (1 mg/ml) in running buffer was flowed over biotinylated gold surface chip at a flow rate of 20 μl/min. The running buffer used was 10 mM HEPES, 150 mM NaCl buffer at pH 7.2 and supplemented with 0.1 mg/ml of BSA and 0.05% (v/v) Tween-20 as recommended by the vendor. ACE2-biotinylated was immobilized on the streptavidin surface by injection of 150 μL at a concentration of 20 μg/ml and at a flow rate of 10 μl/min. Through performing single-cycle kinetics, dissociation times were monitored for at least 900s before each injection of analytes. We used concentrations of 25 nM, 10 nM, 5 nM, and 2.5 nM of RBD-Fc monomers and equivalent RBD concentration with the RBD-PB NPs. RBD and RBD-PB were analyzed in separate experiments. Measurements for each concentration were performed at least in triplicate. Nicoya OpenSPR instrument was used to fit the kinetic data using a 1:1 Langmuir binding model.

#### Size measurement of PB NPs

The hydrodynamic diameter of the different PB NPs used was determined via dynamic light scattering (DLS) with a NanoZetaSizer (Malvern Instruments, Ltd). Independent measurements (N=3) of 100 nM PB were performed in 1X phosphate buffer saline (PBS) at 25°C. Intensity-weighted Z-average diameter for each PB NPs were reported along with their polydispersity index (PDI).

### Animal immunization and viral challenge

#### Immunization assay

We prepared solution of 50 μl of four different PB constructs: PB alone, PB with RBD trimer (1 μg and 5 μg doses), PB with the reconstituted RBD trimer (1 μg dose) along with CpG ODN1018 that were administered to 6 to 8 weeks old female BALB/c mice via intramuscular (IM) injection in the right caudal thigh muscle. Two injections were done with three-week intervals. Animals were examined daily to assess indications of distress according to the parameters defined by ‘Animal Study Clinical Monitoring Chart’ approved by the GMU IACUC. After three weeks from the second injection, animals were euthanized. Serum separated from blood collected from submandibular vein were examined via plaque reduction neutralization assay (PRNT).

#### Plaque reduction neutralization assay (PRNT)

Neutralization antibody titers from each PB nanovaccines were determined through virus inhibition with PRNT assays. Sera collected three weeks after the second injection was two-fold serially diluted in four steps starting with 1:10 dilution in Dulbecco’s Modified Eagle Medium (DMEM) supplemented with 5% fetal bovine serum, 1% L-glutamate, 20 U/ml penicillin, and 20 ug/ml streptomycin. Each sera dilutions were mixed with 100 pfu of virus (SARS-CoV-2, Isolate Italy-INMI1) and serum-virus mix was incubated at 37°C with 5% CO_2_ for 1hr. After incubation, mixture was inoculated into confluent layer of Vero cells in a 12-well plate and incubated at 37°C with 5% CO_2_ for 1 hr. After an hour, 0.6% agarose (ThermoFisher, 16500100) containing Eagles’s Minimum Essential Medium (EMEM– without phenol red) supplemented with 5% FBE, non-essential amino acids, 1 mM sodium pyruvate (VWR, 45000-710, Dixon, CA, USA), 2 mM L-glutamine, 20 U/mL penicillin, 20 μg/mL streptomycin was added to each well in a 1:1 volume ratio. The cells were then incubated at 37°C with 5% CO2 for 48 hrs. After the incubation period, cells were fixed with 10% formaldehyde (Fisher Scientific, F79p-4) for 1 hr. The formaldehyde/agarose plugs were removed after 1 hr. Cells were then washed with deionized water, then stained with 1% crystal violet (FisheSci, C581-25) and 20% ethanol solution (FisherSci, BP2818-4). Plaques were counted and analyzed in a plot showing dilution versus pfu values.

#### Viral inhalational challenge with SARS-CoV-2 strain Isolate Italy-INMI1

6-8-week-old male K18-Ace2 (B6.Cg-Tg(K18-ACE2)2Prlmn/J) mice were subjected to challenge studies with five different PB nanovaccine constructs: PB alone, PB with RBD trimer (1 μg and 5 μg doses), PB with RBD trimer (1 μg dose) and CpG ODN 1018, and PB with CpG ODN1018. 50 μl of each construct was injected via IM route from the right caudal thigh for initial immunization. After three weeks, a second injection was performed. Animals were monitored daily to observe distress parameters in ABSL2 vivarium. At the end of 6^th^ week, animals were moved to ABSL3 vivarium and infected with SARS-CoV-2 (Isolate Italy-INMI1) at a dose of 5×10^4^ pfu through intranasal route. Each animal injected with different constructs were monitored according to individual distress parameters and the survival rate was recorded by ‘Animal Study Clinical Monitoring Chart’ provided by Animal Care Services.

## Ethical Statement

All animal studies carried out for these studies were in accordance with recommendations of the Institutional Animal Care and Use Committee (IACUC protocol #0399) at GMU.

## Statistical Analysis

Stability data of PB nanoparticles were reported as mean and standard deviation as shown on the graph. Due to the small sample size (n:2 for RBD-PB), nanovaccines in stability assays weren’t compared in any statistical test. Data from virus inhibition per diluted sera (12-point dilution) was shown as mean and standard deviation on the graph. Statistical significance was determined via two-way ANOVA, where the p-value was calculated less than 0.001 using R studio version 1.2.5033. Body weight changes (N=5 per study group) of five animals 14 days after challenge was analyzed with one-way ANOVA, Tukey post-hoc test, and Kruskal Wallis post-hoc test. All statistical examinations were performed using R studio.

## Reporting summary

Further information on research design is available in the Nature Research Reporting Summary linked to this article.

## Data availability

All raw and processed data can be made available upon reasonable request to the corresponding authors.

## Acknowledgements

This study was partly funded by the DOD award number W81XWH2010054 and by a COS/VSE Collaborative Seed Fund award. We thank for the support of E.O received by the Turkish Ministry of National Education via the ‘MoNE Scholarship’. We also thank Amanda Graf for her contribution on the production of DNA origami nanoparticle scaffold.

## Author contributions

R.V. conceptualized and designed the *in vitro* studies. A.N., F.A., and R.V. conceptualized and designed the *in vivo* studies. E.O. performed DNA-NP vaccine design, assembly, and characterization experiments. F.A. and K.H. performed the animal experiments. F.A. and K.H. collected animal data. E.O., F.A., and K.H. performed statistical analysis. E.O and R.V wrote the manuscript. All authors reviewed and edited the manuscript.

