## Supplemental Tables 1 to 7 and supplemental Figures 1 to 9 for "DNA Origami Presenting the Receptor Binding Domain of SARS-CoV-2 Elicit Robust Protective Immune Response"

### SUPPLEMENTARY TABLES

**Supplementary Table 1 ssDNA scaffold sequence.** The primer binding sequences are shown in red.

| **PB ssDNA scaffold sequence** |
|---|
| **GCGACGATTTACAGAAGCAA**GGTTATTCACTCACATATATTGATTTATGTACTGTTTCCATTAAAA<br>AAGGTAATTCAAATGAAATTGTTAAATGTAATTAATTTTGTTTTCTTGATGTTTGTTTCATCATCTTC<br>TTTTGCTCAGGTAATTGAAATGAATAATTCGCCTCTGCGCGATTTTGTAACTTGGTATTCAAAGCAA<br>TCAGGCGAATCCGTTATTGTTTCTCCCGATGTAAAAGGTACTGTTACTGTATATTCATCTGACGTTAA<br>ACCTGAAAATCTACGCAATTTCTTTATTTCTGTTTTACGTGCTAATAATTTTGATATGGTTGGTTCAA<br>TTCCTTCCATAATTCAGAAGTATAATCCAAACAATCAGGATTATATTGATGAATTGCCATCATCTGA<br>TAATCAGGAATATGATGATAATTCCGCTCCTTCTGGTGGTTTCTTTGTTCCGCAAAATGATAATGTTA<br>CTCAAACTTTTAAAATTAATAACGTTCGGGCAAAGGATTTAATACGAGTTGTCGAATTGTTTGTAAA<br>GTCTAATACTTCTAAATCCTCAAATGTATTATCTATTGACGGCTCTAATCTATTAGTTGTTAGTGCAC<br>CTAAAGATATTTTAGATAACCTTCCTCAATTCCTTTCTACTGTTGATTTGCCAACTGACCAGATATTG<br>ATTGAGGGTTTGATATTTGAGGTTCAGCAAGGTGATGCTTTAGATTTTTCATTTGCTGCTGGCTCTCA<br>GCGTGGCACTGTTGCAGGCGGTGTTAATACTGACCGCCTCACCTCTGTTTTATCTTCTGCTGGTGGTT<br>CGTTCGGTATTTTTAATGGCGATGTTTTAGGGCTATCAGTTCGCGCATTAAAGACTAATAGCCATTC<br>AAAAATATTGTCTGTGCCACGTATTCTTACGCTTTCAGGTCAGAAGGGTTCTATCTCTGTTGGCCAG<br>AATGTCCCTTTTATTACTGGTCGTGTGACTGGTGAATCTGCCAATGTAAATAATCCATTTCAGACGA<br>TTGAGCGTCAAAATGTAGGTATTTCCATGAGCGTTTTTCCTGTTGCAATGGCTGGCGGTAATATTGTT<br>CTGGATATTACCAGCAAGGCCGATAGTTTGAGTTCTTCTACTCAGGCAAGTGATGTTATTACTAATC<br>AAAGAAGTATTGCTACAACGGTTAATTTGCGTGATGGACAGACTCTTTTACTCGGTGGCCTCACTGA<br>TTATAAAAACACTTCTCAAGATTCTGGCGTACCGTTCCTGTCTAAAATCCCTTTAATCGGCCTCCTGT<br>TTAGCTCCCGCTCTGATTCCAACGAGGAAAGCACGTTATACGTGCTCGTCAAAGCAACCATAGTACG<br>CGCCCTGTAGCGGCGCATTAAGCGCGGCGGGTGTGGTGGTTACGCGCAGCGTGACCGCTACACTTG<br>CCAGCGCCCTAGCGCCCGCTCCTTTCGCTTTCTTCCCTTCCTTTCTCGCCACGTTCGCCGGCTTTCCCC<br>GTCAAGCTCTAAATCGGGGGCTCCCTTTAGGGTTCCGATTTAGTGCTTTACGGCACCTCGACCCCAA<br>AAAACTTGATTTGGGTGATGGTTCACGTAGTGGGCCATCGCCCTG**ATAGACGGTTTTTCGCCCTT** |

**Supplementary Table 2 Primer set for the PB ssDNA scaffold synthesis**

| | |
|---|---|
| PB forward primer | GCGACGATTTACAGAAGCAA |
| PB reverse primer | AAGGGCGAAAAACCGTCTAT |

**Supplementary Table 3 Unmodified staple sequences for folding the PB**

| **Name** | **Sequence** |
|---|---|
| PB-2 | CCACCGAGTAATTTTTAAGAGTCTGTTCTTTGATTAGTTTTTTAATAACATC |

| | |
|---|---|
| PB-3 | GGCCTTGCTGGTTTTTTAATATCCAGTAGACAGGAACTTTTTGGTACGCCAG |
| PB-4 | GAAGGTTATCTTTTTTAAAATATCTTCGTCAATAGATTTTTTAATACATTTG |
| PB-5 | ATATAATCGTTGGCAAATCAACAGTAGAAAG |
| PB-6 | GAATTGAGTATCAGATGATGGCAATTCATCA |
| PB-7 | CGCTGGCAAAGCGAAAGGAGCGGGTATTAAT |
| PB-8 | TTTAAAAGCCTTTGCCCGAACGTCGCTAGGG |
| PB-9 | GCAATACTCCATCACGCAAATTAATAGACTT |
| PB-10 | TACAAACAAGGATTTAGAAGTATCCGTTGTA |
| PB-11 | ATTCGACAACTTTTTTCGTATTAAATTTTGAGTAACATTTTTTTATCATTTTAATTATCATCATTTTTATTCCTGAT |
| PB-12 | AGGAGCGGGCGGAACAAAGAAACATGATGAA |
| PB-13 | ACAAACATACCTGAGCAAAAGAAGCACCAGA |
| PB-14 | ATTAGAGCTAGGTGCACTAACAATAAGAATA |
| PB-15 | CGTGGCACTTCTGACCTGAAAGCGCTAATAG |
| PB-16 | GAGAGCCAAACAGAAATAAAGAAATTGCGTA |
| PB-17 | GATTTTCAACCGCCTGCAACAGTGCCACGCT |
| PB-18 | GGGAGAAACAATATATGTGAGTGGGCCCACT |
| PB-19 | ACGTGAACACAGTAACAGTACCTTTTACATC |
| PB-20 | GGTTTAACGTCTTTTTAGATGAATATCATCACCCAAATTTTTTCAAGTTTTT |
| PB-21 | TCCTCGTTGGATTTTTATCAGAGCGGAAAACGCTCATTTTTTGGAAATACCTGGTGAGGCGGTTTTTTCAGTATTAAC |
| PB-22 | ACGTGCTTTGGGGTCGAGGTGCCGTAAAGCA |
| PB-23 | CTAAATCGGGTTGCTTTGACGAGCACGTATA |
| PB-24 | AAAACAGAACATTTTGACGCTCAATCGTCTG |
| PB-25 | AAATGGATCGAACGAACCACCAGCAGAAGAT |
| PB-26 | TGGCGAGAAAGTTTTTGAAGGGAAGAAGTGTAGCGGTTTTTTCACGCTGCGC |
| PB-27 | CAGTGAGGAATCTTGAGAAGTGTTGCCGCGC |
| PB-28 | CCGCTACAGGGTTTTTCGCGTACTATGAACCCTAAAGTTTTTGGAGCCCCCG |
| PB-29 | TTAATGCGGTAACCACCACACCCTTTATAAT |

| | |
|---|---|
| PB-30 | CCCTCAATCAATTTTTTATCTGGTCACTGATTGTTTGTTTTTGATTATACTT |
| PB-31 | ATCAAAATTATTTTTTTAGCACGTAAGCAGCAAATGATTTTTAAAATCTAAA |
| PB-32 | CGCGCAGAGCTTTGAATACCAAGTAATTGAA |
| PB-33 | CCAACCATCTGAATTATGGAAGGTACAAAAT |
| PB-34 | TAGCCCTAATTAGTCTTTAATGCGACCTCAA |
| PB-35 | ATATCAAAGCATCACCTTGCTGACGAACTGA |
| PB-36 | TATTTACATTGTTTTTGCAGATTCACTGGCCAACAGATTTTTGATAGAACCC |
| PB-37 | AGACAATATTTTTTTTTTGAATGGCTAAACATCGCCATTTTTTTAAAAATAC |
| PB-38 | AAACTATCACTTGCCTGAGTAGAAATAAAAG |
| PB-39 | GGACATTCCAGTCACACGACCAGTAGAACTC |
| PB-40 | GGCGAATTATTTTTTTCATTTCAATTCAAGAAAACAATTTTTAATTAATTAC |
| PB-41 | TAATGGAAACATTTTTGTACATAAATCAATAACGGATTTTTTTCGCCTGATT |
| PB-42 | GGCGAACGATTTAGAGCTTGACGGTTGAATT |
| PB-43 | ACCTTTTTATTTAACAATTTCATGGAAAGCC |
| PB-44 | AGGGATTTAACAATATTACCGCCAGCCATTG |
| PB-45 | CAACAGGAGAGCTAAACAGGAGGCCGATTAA |

**Supplementary Table 4 Modified staples for Antigen-PNA binding.** Overhang sequences are underlined.

| **Name** | **Sequence** |
|---|---|
| PB-17 w/overhang | GATTTTCAACCGCCTGCAACAGTGCCACGCTTTACTGGACTG |
| PB-19 w/overhang | ACGTGAACACAGTAACAGTACCTTTTACATCTTACTGGACTG |
| PB-22 w/overhang | ACGTGCTTTGGGGTCGAGGTGCCGTAAAGCATTACTGGACTG |
| PB-25 w/overhang | AAATGGATCGAACGAACCACCAGCAGAAGATTTACTGGACTG |
| PB-27 w/overhang | CAGTGAGGAATCTTGAGAAGTGTTGCCGCGCTTACTGGACTG |
| PB-32 w/overhang | CGCGCAGAGCTTTGAATACCAAGTAATTGAATTACTGGACTG |

| | |
|---|---|
| PB-35 w/overhang | ATATCAAAGCATCACCTTGCTGACGAACTGATTACTGGACTG |
| PB-38 w/overhang | AAACTATCACTTGCCTGAGTAGAAATAAAAGTTACTGGACTG |
| PB-42 w/overhang | GGCGAACGATTTAGAGCTTGACGGTTGAATTTTACTGGACTG |
| PB-45 w/overhang | CAACAGGAGAGCTAAACAGGAGGCCGATTAATTACTGGACTG |

### Supplementary Table 5 Modified staples with overhang to hybridize CpG sequences.

Overhang sequences are underlined.

| **Name** | **Sequence** |
|---|---|
| PB-5 w/overhang | ATATAATCGTTGGCAAATCAACAGTAGAAAGTAAGTTAGGACCATGAAGT |
| PB-6 w/overhang | GAATTGAGTATCAGATGATGGCAATTCATCATAAGTTAGGACCATGAAGT |
| PB-7 w/overhang | CGCTGGCAAAGCGAAAGGAGCGGGTATTAATTAAGTTAGGACCATGAAGT |
| PB-8 w/overhang | TTTAAAAGCCTTTGCCCGAACGTCGCTAGGGTAAGTTAGGACCATGAAGT |
| PB-9 w/overhang | GCAATACTCCATCACGCAAATTAATAGACTTTAAGTTAGGACCATGAAGT |
| PB-10 w/overhang | TACAAACAAGGATTTAGAAGTATCCGTTGTATAAGTTAGGACCATGAAGT |
| PB-12 w/overhang | AGGAGCGGGCGGAACAAAGAAACATGATGAATAAGTTAGGACCATGAAGT |
| PB-13 w/overhang | ACAAACATACCTGAGCAAAAGAAGCACCAGATAAGTTAGGACCATGAAGT |
| PB-14 w/overhang | ATTAGAGCTAGGTGCACTAACAATAAGAATATAAGTTAGGACCATGAAGT |
| PB-15 w/overhang | CGTGGCACTTCTGACCTGAAAGCGCTAATAGTAAGTTAGGACCATGAAGT |

### Supplementary Table 6 FRET-dye modified staple strands

| **Name** | **Sequence** |
|---|---|
| Fret1-Don1 | FAM//ATTCGACAACTTTTTTCGTATTAAATTTTGAGTAACATTTTTTTATCATTTTAATTATCATCATTTTTATTCCTGAT |
| Fret1-Acc1 | TAMRA//TACAAACAAGGATTTAGAAGTATCCGTTGTA |

| | |
|---|---|
| Fret2-Don2 | FAM//TCCTCGTTGGATTTTTATCAGAGCGGAAAACGCTCATTTTTTGGAAATACCTGGTGAGGCGGTTTTTTCAGTATTAAC |
| Fret2-Acc2 | TAMRA//ACGTGCTTTGGGGTCGAGGTGCCGTAAAGCA |

**Supplementary Table 7 Examples of Kd extracted from the literature for SARS-CoV-2 RBD binding to the human ACE2 based on BLI and SPR results.**

| **Receptor-Ligand** | **Kd (nM)** |
|---|---|
| **ACE2-RBD** | 4.7 [1] |
| **ACE2-RBD** | 3.2 [2] |
| **ACE2-RBD, Fc** | 7.16 [2] |
| **ACE2-RBD dimer** | 1.25 [2] |
| **ACE2-RBD** | 1.59 [3] |
| **ACE2-RBD** | 19.3[4] |
| **ACE2-RBD trimer** | 0.04[4] |

**SUPPLEMENTARY FIGURES**

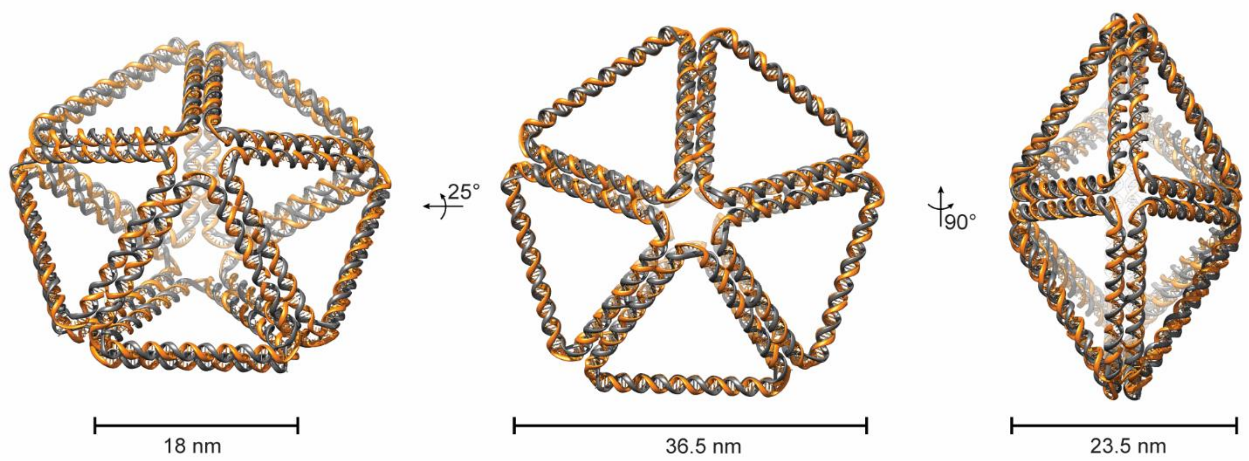


**Supplementary Fig. 1 Design of DNA pentagonal bipyramid (PB)**. PB was designed with DAEDALUS and the rendering performed with Chimera.

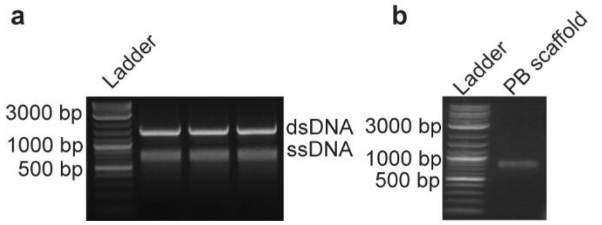


**Supplementary Fig. 2 PB ssDNA scaffold production.** The 1,616 nts ssDNA scaffold was synthetized with aPCR. **a** Agarose gel electrophoresis showing the results of three independent aPCR reactions. **b** Validation of the ssDNA scaffold purity.

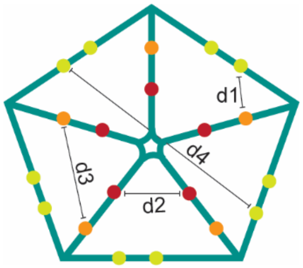


**Supplementary Fig. 3 Graphical representation of the distances between the antigen-binding sites on the PB NP**. (d: distance) d1= 9 nm; d2= 11 nm; d3= 15 nm; d4= 36.5 nm. Distances were measured in Chimera using the model generated by DAEDALUS.

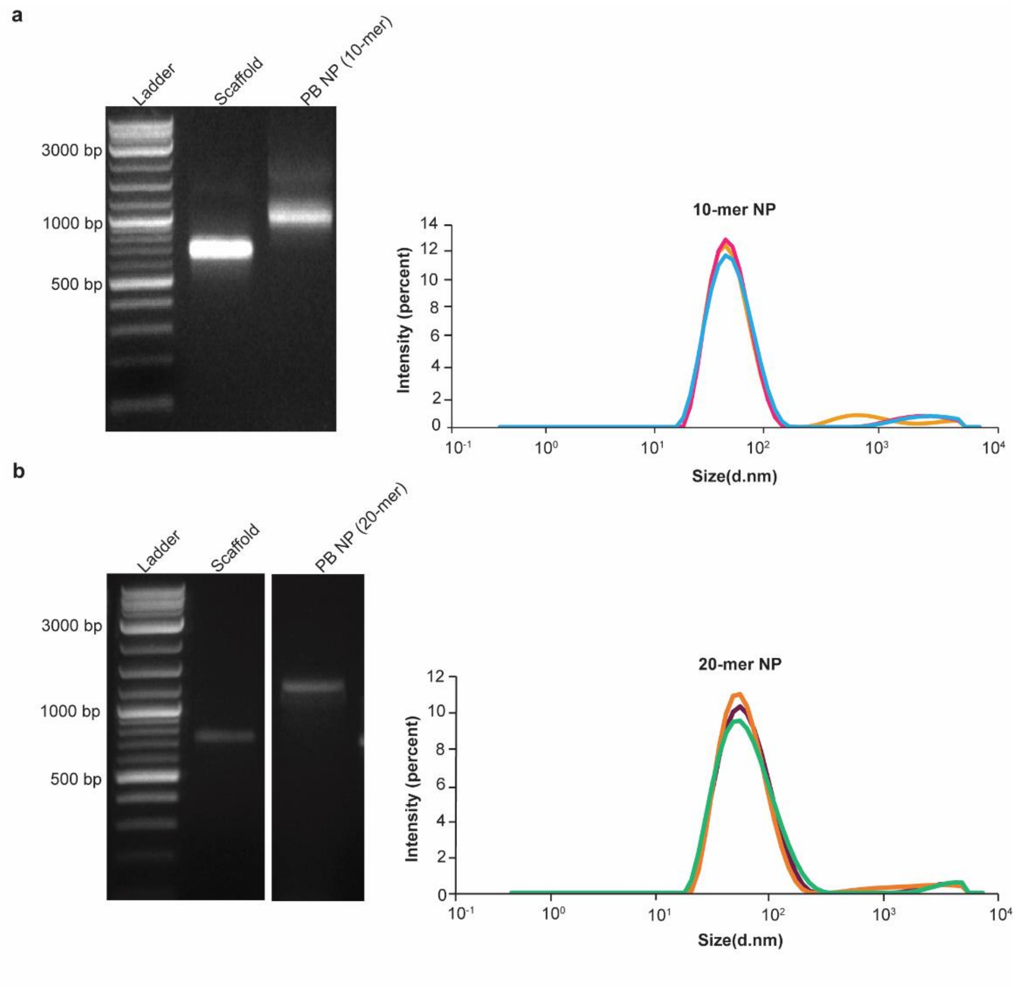


**Supplementary Fig. 4 a** Characterization of PB-10 (10 overhangs on one face of PB) via agarose gel electrophoresis and DLS. Hydrodynamic diameter (Z-average= 47.4 nm) measured from three different NP preparations. **b** Characterization of PB-20 (10 overhangs on each face of PB) via agarose gel electrophoresis and DLS. Hydrodynamic diameter (Z-average= 51.4 nm) measured from three different NP preparations. Theoretical diameters were estimated at 40.2 nm and 46.4 nm for PB-10 and PB-20, respectively. Polydispersity index measured for PB-10 is 0.241 and for PB-20 is 0.243.

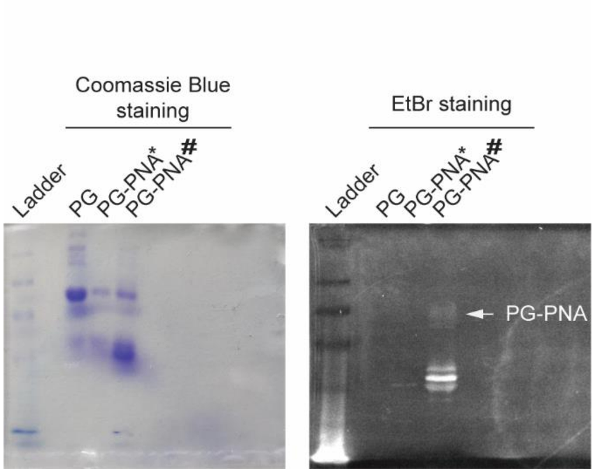


**Supplementary Fig. 5 Validation of PNA-maleimide conjugated PG in denaturing gel.** 10% SDS-PAGE (sodium dodecyl sulfate polyacrylamide gel electrophoresis) gel was initially stained with PageBlue protein staining solution to visualize PG. Then the same gel was incubated with EtBr to visualize PNA-conjugated PG. White arrow indicates PG-PNA. Approximate molecular mass of PG is 21.9 kDA and migrate in SDS-PAGE with molecular weight of 40 kDA. Theoretical mass of PG-PNA is 24.9 kDA. * indicate that the PG was reacted with PNA without TCEP reduction, # indicate the PG-PNA was first reduced with TCEP and then reacted with PNA overnight.

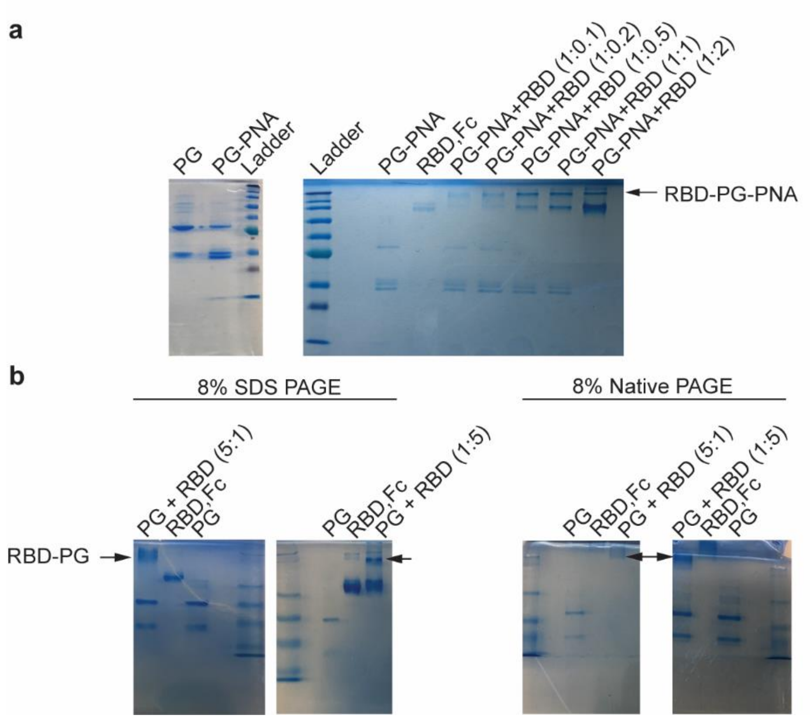


**Supplementary Fig. 6 Formation of the RBD trimer. a** PG-PNA is incubated with different molar ratios of RBD-Fc. The conjugation of two proteins was evaluated through running the samples in SDS-PAGE gel. **b** Additionally, five-fold excess of PG without PNA was incubated with RBD-Fc, and five-fold excess of RBD-Fc was incubated with PG without PNA. Conjugation of two proteins was examined via native and denaturing gel electrophoresis. Theoretical molecular mass of RBD, Fc is 51.5 kDA and migrate in SDS PAGE approximately between 60 and 65 kDA. Theoretical mass of RBD-PG is 176.4 kDA and RBD-PG-PNA is 179.4 kDA.

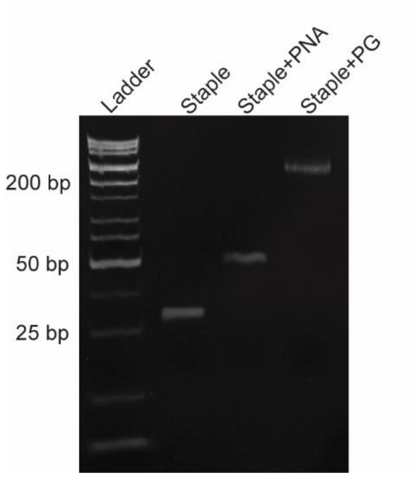


**Supplementary Fig. 7 Protein G-PNA binding to the DNA overhangs.** An individual staple strand with an overhang was incubated with PNA and PG-PNA at 37°C. We used 14% PAGE gel to evaluate the hybridization of PNA alone and PG-PNA to the overhang.

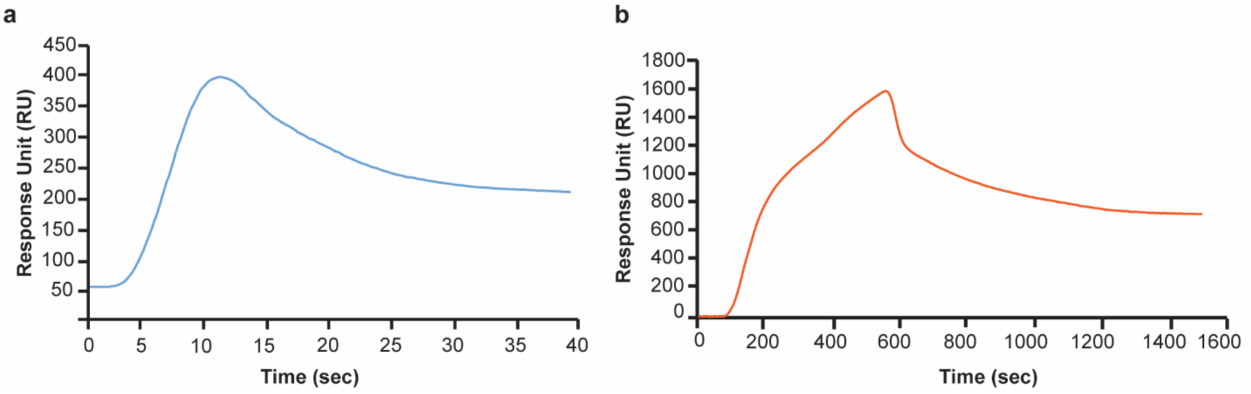


**Supplementary Fig. 8 Representative SPR binding curve**. **a** Streptavidin binding on biotinylated gold sensor. **b** Immobilization of the ACE2-biotinylated receptor on streptavidin-modified gold sensor.

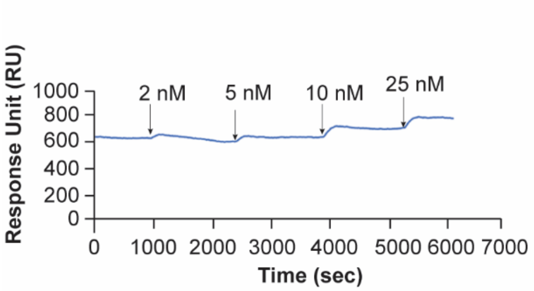


**Supplementary Fig. 9 Sequential binding interactions of RBD with the ACE2 receptor for four different concentrations of RBD-PB NP during single cycle kinetics (SCK) experiment.**
